# Comparative Molecular Dynamics Characterization of Hair Keratin Unfolding Mechanics^*^

**DOI:** 10.64898/2026.06.06.730563

**Authors:** Wei Lu, Fabien Leonforte, Markus J. Buehler

## Abstract

Keratin proteins are fundamental structural components of hair fibers, contributing to their mechanical resilience, elasticity, and fracture resistance. However, systematic molecular-scale characterization of keratin unfolding mechanics across protein types remains limited, restricting the connection between protein-level deformation mechanisms and hierarchical hair fiber mechanics. Here, we establish a comparative molecular-dynamics-based framework for characterizing the unfolding behavior and nanomechanical response of a curated dataset of 51 keratin proteins. We conduct implicit atomistic molecular dynamics (MD) simulations, including equilibration and steered molecular dynamics (SMD) under four accelerated pulling velocities, to quantify unfolding forces, energy absorption, and structure–property relationships. These accelerated pulling conditions are interpreted as computational probes of relative molecular-scale trends, rather than direct reproductions of experimental hair-fiber strain-rate regimes. Across these accelerated SMD conditions, the simulations show rate-sensitive increases in unfolding force and energy absorption, consistent with constrained molecular relaxation during faster molecular pulling. Stronger correlations between nanomechanical properties and molecular descriptors emerge at higher pulling rates, and the nanomechanical responses of different keratin types (Type I and II) are also compared. The findings provide molecular-level insights into protein unfolding mechanisms that may contribute to the mechanical behavior of hierarchical keratin structures. This study establishes a quantitative framework for comparative keratin unfolding mechanics, providing molecular-level descriptors for future multiscale modeling of hair fiber behavior. These results support applications in biomaterial design, hair fiber durability analysis, and bioinspired material engineering. Future work will integrate these nanomechanical descriptors with fiber-level mechanics and machine learning-based keratin design.

## 1 Introduction

Human hair fibers, as a biological material, primarily composed of keratin protein, exhibiting high strength [1] and elasticity [2], featuring hierarchical structures, ranging from nanoscale amino acid chains to microscale hair fibers [3, 4] as visualized in Figure 1(a). At the nanoscale, amino acids assemble into alpha-helix chains, which form heterodimers composed of both type I and II keratin monomers. These heterodimers further organize into intermediate filaments, which are built hierarchically from tetramers to protofilaments and then to protofibrils. At the next scale, macrofibrils are formed, which combine to create cortical cells and, ultimately, microscale hair fibers (20–100 µm in diameter) composed of the cortex, medulla, and cuticle layers. For hair fiber layers [3, 5], cortex layer is crucial for determining the mechanical properties of hair fibers such as strength, and outermost cuticle layer protects hair fiber and maintains its integrity, and medulla, a disordered region can be observed in the central region of some fibers. The hair cycle can be described in three stages [6, 7]: anagen (growth), catagen (regression), and telogen (rest). During anagen, hair follicles actively produce a new hair shaft, extending from tip to root. In catagen, the follicle undergoes regression, resetting its structure, while telogen serves as a quiescent phase, preparing stem cells for the next growth cycle.

**Figure 1.**
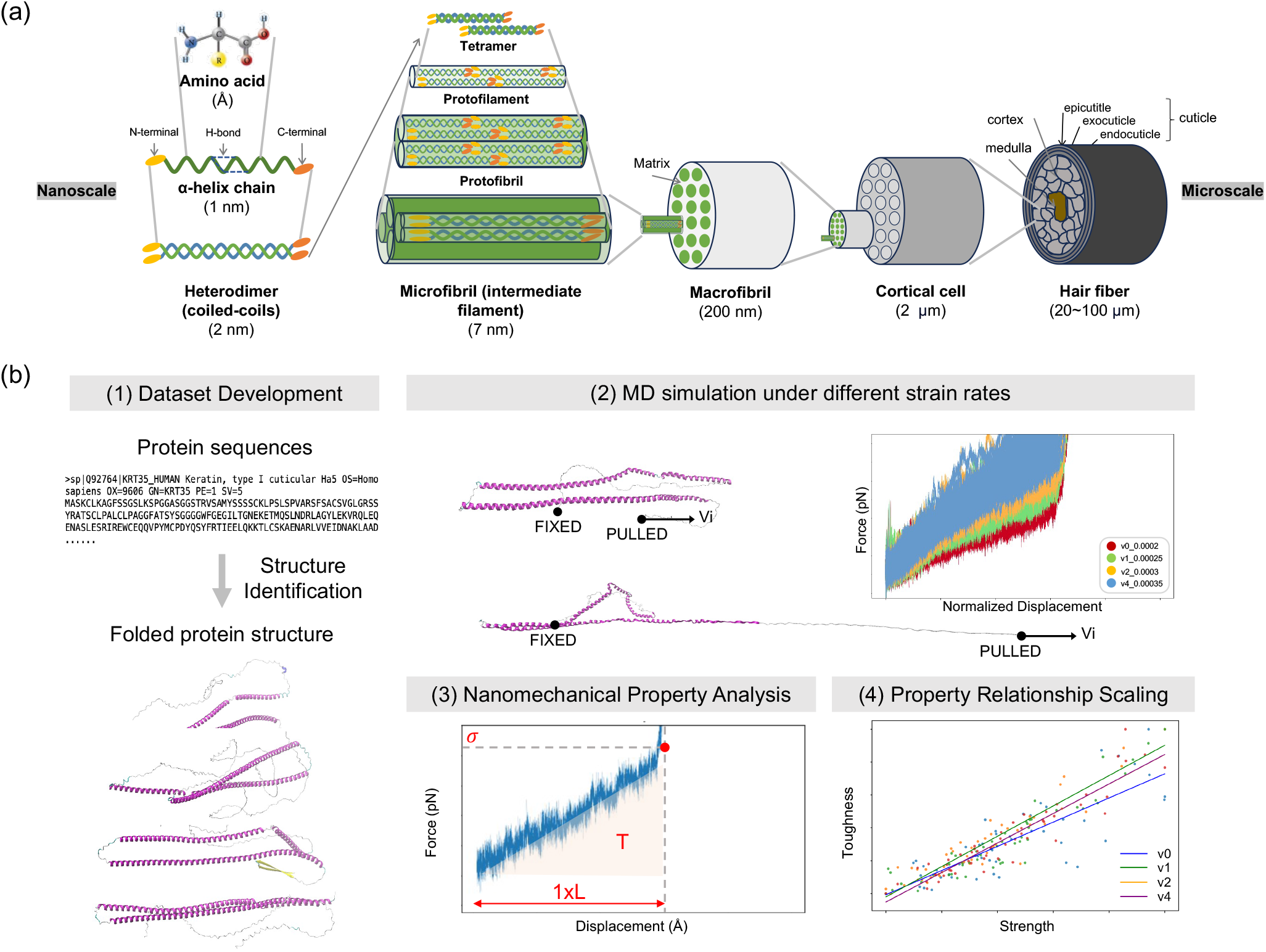
Schematic of the hierarchical structure of a hair fiber and overview of the project workflow. Panel (a) shows the hierarchical structure of a hair fiber. At the nanoscale, amino acids assemble into alpha-helix chains, which form heterodimers composed of both type I and II keratin monomers. These heterodimers further organize into intermediate filaments, which are built hierarchically from tetramers to protofilaments and then to protofibrils. At the next scale, macrofibrils are formed, which combine to create cortical cells and, ultimately, microscale hair fibers (20–100 µm in diameter) composed of the cortex, medulla, and cuticle layers. Panel (b) shows the overall workflow of the project. The process begins with keratin protein dataset development, including keratin protein sequence collection and structure prediction using folding tools. Molecular dynamics (MD) simulations, comprising equilibration and steered molecular dynamics (SMD), are performed to simulate protein stretching under four controlled accelerated pulling velocities. Simulation results are used to characterize nanomechanical properties and structure–property relationships across the keratin dataset.

Hair fibers exhibit distinct physical properties that impact durability, manageability, and styling [8, 9]. When wet, hair can stretch up to 30% of its original length without damage, but beyond 70%, irreversible changes occur, and at 80%, fracture happens. Porosity affects water absorption, with damaged hair being highly porous and more prone to breakage due to increased combing friction when wet. Dry hair generates static electricity, causing flyaways, which moisture can reduce. Human hair is primarily classified by structure and ethnicity[10, 11]. Structurally, hair varies from straight to wavy, curly, and kinky, influencing texture and porosity. Ethnic variations include Asian hair (straight, thick), Caucasian hair (diverse textures), and African hair (coiled, fragile), each exhibiting distinct mechanical behaviors[12, 13]. Asian and Caucasian hair generally have higher tensile strength, while African hair, with its kinks and coils, is more fragile and prone to breakage. Curly hair displays a toe region in stress-strain curves, indicating initial resistance before extension. Bending elasticity increases with curliness, while thicker Asian hair exhibits higher torsional stiffness. Surface roughness is greater in Caucasian and African hair, while Asian hair is more resistant to hydration changes. UV and chemical resistance also vary, with darker, curlier hair retaining proteins better but experiencing greater lipid degradation, influencing hair durability, treatment responses, and product formulations [12].

The strength and stability of hair fibers depend on three key intermolecular interactions: disulfide bonds, Coulomb interactions, and hydrogen bonds [14, 15, 16]. Disulfide bonds, formed by cystine cross-links, provide structural integrity but are highly susceptible to oxidation and reduction, making them a primary target in chemical treatments. Coulomb interactions, arising from acidic and basic side chains, contribute to fiber stability in aqueous environments but break under acidic or alkaline conditions. Hydrogen bonds, though weak and water-sensitive, are the most abundant and crucial for maintaining keratin’s alpha-helical structure. Together, these intermolecular interactions define the mechanical resilience and chemical behavior of hair fibers, shaping their response to treatments, environmental exposure, and grooming practices.

When considering the protein-scale composition, keratin as an intermediate filament (KIF) protein is the key component. It can be categorized based on their isoelectric point (pI): acidic type I keratin with pI of about 4.9 to 5.4, and basic or neutral type II keratin with pI of about 6.5 to 8.5 [17]. These two types combine to form a heterodimer, aligning in parallel and contributing to the overall structure of hair [17]. In addition to keratin, hair keratin-associated proteins (KAP) represent another essential component, play integral roles in establishing the hair shaft’s strength by forming cross-links with keratin intermediate filaments (KIF) [18, 19]. KAPs can be classified into three categories based on amino acid composition [20], specifically the sulfur content and glycine-tyrosine levels, which influence their structural and functional roles in hair fiber formation: high-sulfur proteins, ultra-high-sulfur proteins, and glycine-tyrosine-rich proteins. The interaction of keratins with keratin-associated proteins (KAPs) further reinforces the hair fiber matrix, affecting its mechanical and chemical properties.

Hair issues arise from mechanical, chemical, environmental, and biological factors, impacting hair health and strength[7]. Mechanical damage from brushing, tight hairstyles, and heat styling leads to split ends and white dots, while chemical treatments like bleaching and perming weaken keratin, making hair brittle and porous. Environmental stressors such as UV radiation, pollution, and humidity accelerate cuticle damage and dehydration, and scalp conditions like dandruff cause flakiness and inflammation, contributing to hair thinning. Biological factors, including hormonal imbalances, stress, and medical conditions, result in hair loss and premature graying. Maintaining gentle hair care, minimizing damage, and supporting scalp health are crucial for hair longevity[7, 10, 8]. Moisturizing shampoos, conditioners, and deep treatments help restore moisture and elasticity, while protective hairstyles and reduced heat exposure prevent breakage, especially in curly and coiled hair. Avoiding tight hairstyles, excessive chemical treatments, and stress helps prevent hair loss, while advances in conditioning polymers, protein treatments, and UV protection enhance resilience, shine, and manageability, counteracting daily stressors.

Extensive research has been conducted on hair fibers and keratin proteins, employing various experimental and computational approaches. Studies have explored mechanical properties of hair, such as splitting behavior through comparative analysis [24], structural mechanisms of hair fibers [1, 25], and hair fiber composites [26]. Chemical and environmental impact studies have investigated hair bleaching effects [27] and photochemical degradation [28]. Additionally, microscopy-based morphological analyses using SEM and TEM have provided insights into hair fiber microstructures and dam-cause flakiness and inflammation, contributing to hair thinning. Biological factors, including hormonal imbalances, stress, and medical conditions, result in hair loss and premature graying. Maintaining gentle hair care, minimizing damage, and supporting scalp health are crucial for hair longevity[7, 10, 8]. Moisturizing shampoos, conditioners, and deep treatments help restore moisture and elasticity, while protective hairstyles and reduced heat exposure prevent breakage, especially in curly and coiled hair. Avoiding tight hairstyles, excessive chemical treatments, and stress helps prevent hair loss, while advances in conditioning polymers, protein treatments, and UV protection enhance resilience, shine, and manageability, counteracting daily stressors.

Extensive research has been conducted on hair fibers and keratin proteins, employing various experimental and computational approaches. Studies have explored mechanical properties of hair, such as splitting behavior through comparative analysis [24], structural mechanisms of hair fibers [1, 25], and hair fiber composites [26]. Chemical and environmental impact studies have investigated hair bleaching effects [27] and photochemical degradation [28]. Additionally, microscopy-based morphological analyses using SEM and TEM have provided insights into hair fiber microstructures and dam-age assessment [29, 30]. Biophysical and biological investigations have examined hair protein chemistry [31, 32], while cosmetic and hair care product evaluations focus on treatment effects and formulations [33, 34, 35]. Finite element modeling (FEM) has been used to analyze hair fiber mechanical responses and fiber-reinforced composite structures [36, 37, 38]. Advanced techniques, including machine learning and emerging agent-based AI frameworks, have been applied to study biopolymer properties in hair [39], detect scalp fungal infections [40], support autonomous scientific discovery workflows [41], and generate synthetic spider silks and bio-inspired materials [42, 43, 44, 45]. Molecular dynamics (MD) simulations have been utilized to investigate keratin interactions [46], the role of disulfide bonds [47], fatty acid interactions in hair fibers [48], and protein unfolding in hierarchical materials [49]. Tools like AlphaFold [50] and OmegaFold [51] have further facilitated protein folding predictions, aiding MD simulations. Despite advancements in hair fiber research, a systematic molecular-scale framework for comparing keratin unfolding mechanics across protein types remains underdeveloped. Most studies focus on static mechanical properties, fiber-level analyses, or specific molecular interactions, while dataset-scale characterization of keratin unfolding, energy dissipation, and structure–property relationships remains limited. Understanding how keratin proteins unfold and dissipate energy under controlled molecular pulling is crucial for linking molecular-scale mechanics to features relevant for fiber-level mechanical performance.

### 1.1 Motivations

Keratin proteins are key structural components of hair fibers, providing mechanical resilience, elasticity, and fracture resistance. While prior research has examined keratin composition, molecular structure, and fiber-level properties, systematic molecular-scale characterization of keratin unfolding mechanics across different keratin proteins remains limited. In particular, the relationships between nanomechanical properties, such as unfolding force and energy absorption, and molecular descriptors, such as sequence length, secondary structure, molecular weight, and solvent accessibility, remain unclear. Addressing these gaps can provide insight into how protein-level deformation mechanisms contribute to hierarchical hair fiber mechanics, damage resistance, and future keratin-based biomaterial design.

It is important to note that the pulling velocities used in steered molecular dynamics (SMD) are far above regular experimental hair-fiber deformation rates. Therefore, this study does not aim to directly reproduce macroscopic hair-fiber deformation under experimentally accessible strain-rate conditions. Instead, the different pulling velocities are used as controlled computational perturbations to probe relative unfolding behavior, resistance to extension, energy dissipation, and structure-property trends across keratin proteins.

Thus, we propose this study to establish a quantitative MD framework for comparative characterization of keratin unfolding mechanics. As illustrated in Figure 1(b), this work develops a curated keratin protein dataset, performs molecular simulations under controlled pulling conditions, quantifies nanomechanical descriptors such as strength and toughness, and examines structure-property relationships that influence unfolding resistance and energy dissipation. By providing a detailed molecular perspective, this study lays the groundwork for multiscale modeling and could guide future efforts in hair fiber mechanics, biomaterial engineering, and synthetic keratin design. Future research could further extend this framework by integrating machine learning techniques to predict keratin mechanical behavior, explore nonlinear correlations, and design synthetic keratin sequences with targeted mechanical properties.

### 1.2 Significance of this work

This study establishes a systematic computational framework to compare molecular unfolding and nanomechanical descriptors across a curated set of hair keratin proteins. While previous studies have examined hair fibers experimentally or explored specific keratin-related mechanisms, dataset-scale molecular analysis of keratin unfolding mechanics remains limited. By combining curated sequence collection, structure prediction, consistent MD simulation protocols, and comparative nanomechanical analysis across 51 keratin proteins, this work offers a quantitative view of keratin unfolding behavior, energy dissipation, and structure–property relationships. The computational workflow also makes the analysis more efficient, reproducible, and scalable, allowing comparison across multiple proteins and loading conditions in a more systematic way.

The novelty of this study lies primarily in its dataset-scale and reproducible comparison of keratin molecular unfolding mechanics, with the applied pulling velocities used as controlled computational perturbations rather than experimentally matched strain-rate conditions. The framework enables quantitative characterization of strength and toughness under controlled SMD conditions and allows their relationships with molecular descriptors, including molecular weight, coil content, and solvent-accessible surface area, to be analyzed in a consistent manner. In addition, the study compares the nanomechanical responses of different keratin classes, specifically Type I and Type II keratins, revealing both shared scaling trends and class-dependent differences in strength and toughness distributions.

More broadly, this framework helps connect molecular unfolding mechanisms to higher-level hair fiber behavior. It also supports future work in multiscale modeling, predictive modeling of keratin mechan-ics, and the design of keratin-based biomaterials or hair treatment strategies.

### 1.3 Paper outline

This paper is organized as follows. We begin with the background of this work, outlining the research motivations, existing gaps in current topics, and the objectives of our study. Next, we present a detailed analysis and discussion of the developed dataset, simulation results, and the characterization of nanomechanical properties and correlations. A comprehensive conclusion summarizes the study, highlighting key findings, potential impacts, and future research directions. Finally, the specific methodologies and tools used for dataset development, simulation, and result analysis are described.

## 2 Results and Discussion

In this section, we present a detailed analysis and discussion of the curated dataset, the simulation performance under controlled SMD pulling velocities, the nanomechanical properties characterized from simulations, and the correlation and scaling of these properties with molecular descriptors. The specific methodologies implemented follow the process outlined in Section 4.

### 2.1 Dataset development

A dataset of 51 keratin proteins was curated for this study, with an average sequence length of 506 residues. Although the availability of open-source keratin protein data is limited, leading to a relatively small dataset size, the quality and structural uncertainties of the protein data were systematically evaluated through property analysis, molecular folding performance, and structural assessment. To further examine structural variations and folding accuracy, seven representative keratin proteins were selected, including four Type I keratins (KRT31, KRT32, KRT35, KRT37) and three Type II keratins (KRT81, KRT85, KRT86). These proteins were chosen based on their classification as Type I (acidic) or Type II (neutral/basic) keratins and their expression sites, predominantly in the cortex, which plays a key role in governing the mechanical properties of hair fibers, as opposed to keratins found in the cuticle or medulla [55, 56]. All 51 collected data is summarized in a CSV file, provided in the supplementary materials, along with calculated protein properties. Additionally, the FASTA protein sequences and folded PDB structures are also included in the supplementary materials.

The sequence-based protein properties are calculated using ProtParam in python [22]. The property distributions are indicated in Figure 2(a) for the 51 keratin proteins in the dataset, which provides insights into their structural and functional characteristics. The sequence length ranges between 400 and 600 residues, with most proteins clustering around 450–550 residues. This relatively narrow distribution suggests a conserved domain organization, which is expected for keratin proteins that play structural roles in intermediate filaments. The molecular weight (MW) distribution spans 45,000 to 65,000 Da, with a peak in the 50,000–60,000 range, indicating that most keratins in this dataset share a similar molecular size. This observation is consistent with keratin proteins’ known mechanical and structural functions, where a specific size range is necessary for proper filament assembly. The instability index values are distributed mostly between 40 and 65, with a peak around 55, suggesting that most keratin proteins are moderately stable. Some proteins exhibit higher II values, indicating the presence of flexible or disordered regions, which may be functionally significant in keratin filament interactions. The isoelectric point distribution exhibits a bimodal pattern, with peaks around 5.0–5.5 and 7.0–8.0, corresponding to Type I (acidic) and Type II (basic/neutral) keratins, respectively. The prominence of the acidic peak (5.0) suggests a higher representation of Type I keratins in the dataset, which aligns with their abundance in certain epithelial tissues. The bimodal nature of the distribution confirms the complementary roles of Type I and Type II keratins in forming heteropolymeric filaments, a crucial feature for keratin fiber stability. Overall, the protein property distributions in this dataset reflect the structural characteristics of keratin proteins. The conserved sequence length and molecular weight support their role in forming stable intermediate filaments, while the instability index values indicate potential flexibility or disorder in some keratins. The bimodal IsoPt distribution reinforces the functional division between Type I and Type II keratins and suggests that the dataset comprises two biologically distinct sub-populations that play complementary roles in filament formation and mechanical stability. Subsequent stratified analysis indicates that these two classes differ in the distributions of nanomechanical properties, while preserving a similar strengthtoughness scaling relationship.

**Figure 2.**
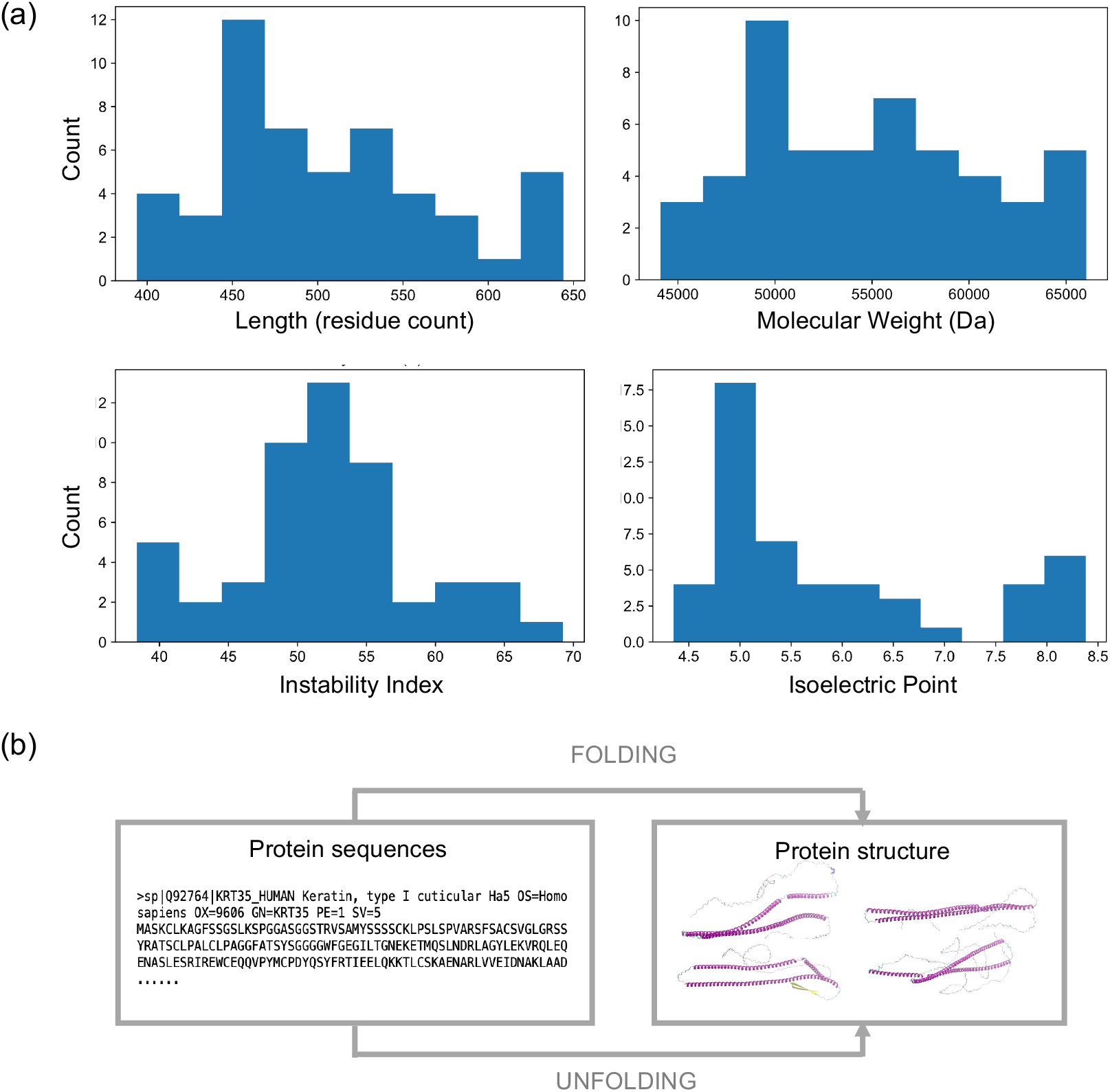
Statistical distribution of keratin protein properties. (a) Protein sequences for 51 keratin proteins, including both Type I (acidic) and Type II (basic/neutral) keratins, were collected from UniProt [21]. Sequence properties were analyzed using ProtParam [22], including length, molecular weight (MW), instability index, and isoelectric point. Length is reported as residue count, MW is reported in Da, and instability index and isoelectric point are dimensionless quantities. The instability index, where values < 40 indicate instability [23], provides insight into protein stability, while IsoPt represents the pH at which the net charge is zero. The distributions show that protein length, MW, and II are uniformly distributed, whereas IsoPt exhibits a bimodal distribution, reflecting the distinction between acidic (Type I) and basic (Type II) keratins. (b) Workflow for protein sequence processing and structure generation, illustrating the transformation from amino acid sequences to folded protein structures used in subsequent simulations.

The molecular structures are folded using AlphaFold2 [50] (Figure 2(a)) based on the collected protein sequences, and the protein structures are visualized using Visual Molecular Dynamics (VMD). The seven sample protein structures and folding prediction performances are illustrated in Figure 3. The molecular structures presented in the figure exhibit the characteristic coiled-coil alpha-helical conformation of keratin proteins. The top row (KRT31, KRT32, KRT35, KRT37) represents Type I keratins, while the bottom row (KRT81, KRT85, KRT86) corresponds to Type II keratins. The structural models display well-defined helical domains (highlighted in purple), which are essential for the mechanical integrity of keratin fibers. However, noticeable structural variations occur in the terminal regions, where extended loops and disordered segments appear, indicating potential flexibility or dynamic interactions. These regions are less structured and likely contribute to protein-protein interactions or adaptability in different environments. The corresponding predicted local distance difference test (pLDDT) plots provide insights into the confidence levels of these structural predictions. The y-axis represents the pLDDT score, with higher values indicating greater confidence in structural accuracy, while the x-axis maps the residue positions along the protein sequence. The results show that the central alpha-helical regions have high pLDDT scores (around over 90), signifying strong structural confidence, while random coils are the N/C-terminis exhibit lower scores (around 30–50), suggesting increased flexibility or disorder. The pattern is observed in most keratins, reflecting the similar molecular structural characteristic of these proteins. The bimodal pLDDT distribution, with high-confidence helical domains and low-confidence termini, corresponds well to the modular architecture of keratin proteins. The structured helices provide mechanical strength, while the flexible termini may play a role in network formation and interactions with other cellular components. Overall, an average pLDDT value of 73.74 was calculated for the 51 selected folded keratin protein structures, indicating good folding prediction quality suitable for further molecular simulations [53, 54].

**Figure 3.**
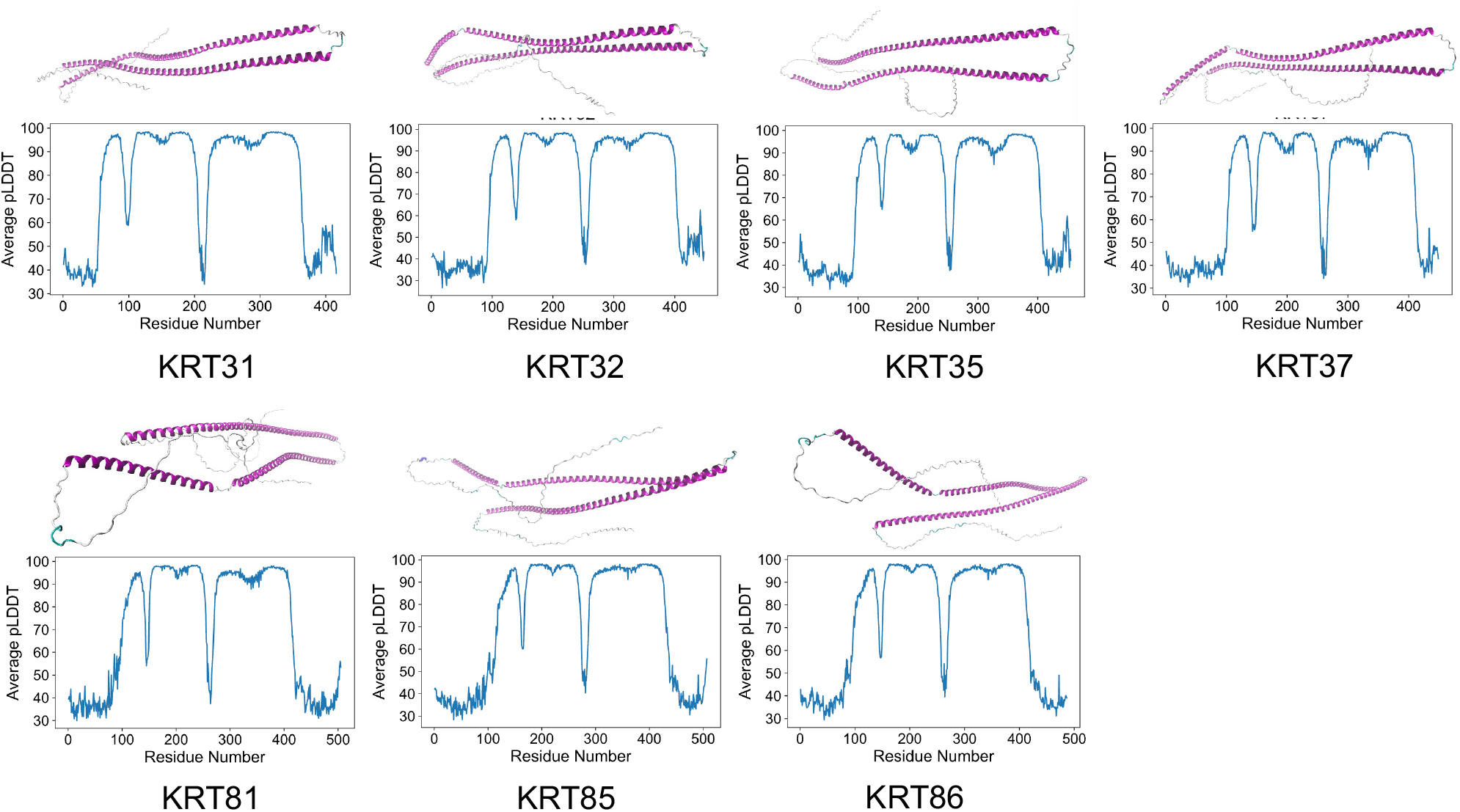
Representative keratin protein structures and folding confidence. Seven representative keratin proteins were selected for folding performance evaluation, and their structures were visualized using Visual Molecular Dynamics (VMD) [52]. The first row represents Type I keratins (KRT31, KRT32, KRT35, KRT37), while the second row represents Type II keratins (KRT81, KRT85, KRT86). The molecular structures presented in the figure exhibit the characteristic coiled-coil alpha-helical conformation of keratin proteins. The structural models display well-defined helical domains (highlighted in purple), primarily connected by turns, with noticeable structural variations in the random coils at the terminal regions. The molecular structures of these keratins exhibit similar overall characteristics, reinforcing their conserved secondary structure organization. The predicted local distance difference test (pLDDT) scores per amino acid from AlphaFold2 [50] are provided for these proteins. An average pLDDT value of 73.74 was obtained across the dataset, with 51 folding pLDDT values calculated, indicating high-quality folding predictions for keratin proteins [53, 54]. Each pLDDT plot illustrates the confidence level of the predicted structures. The y-axis represents the pLDDT score, where higher values indicate greater confidence in structural accuracy, while the x-axis maps residue positions along the protein sequence. A bimodal pLDDT distribution is observed across most keratins, where central alpha-helical regions exhibit high pLDDT scores (consistently over 90), indicating well-structured regions, whereas random coils at the N- and C-termini display lower scores (30–50), reflecting higher flexibility or disorder.

### 2.2 Molecular dynamics simulation performance under controlled SMD pulling velocities

The results of MD simulations, including equilibration and SMD, are presented in Figure 4. The RMSD evolution, force-displacement curves under different SMD pulling velocities, and molecular structure visualizations for example type I protein KRT35 and type II protein KRT85 are shown in panels (a) and (b), respectively. Panel (c) provides a comprehensive view of RMSD trends across all 51 proteins and their collective force-displacement responses under different pulling velocities. Additionally, force distribution comparisons for different SMD pulling velocities are presented in the final plot of panel (c). The RMSD plots in panels (a) and (c) show expected fluctuations during equilibration, which stabilize after approximately 400 ps, indicating that the protein structures have reached a quasiequilibrium state before SMD. However, variations persist among individual equilibration runs due to factors such as inherent thermal fluctuations, differences in initial velocity assignments, and solvent approximations in the implicit GBIS model. These variations in RMSD, especially in implicit solvent models, lead to slightly different equilibrated conformations.

**Figure 4.**
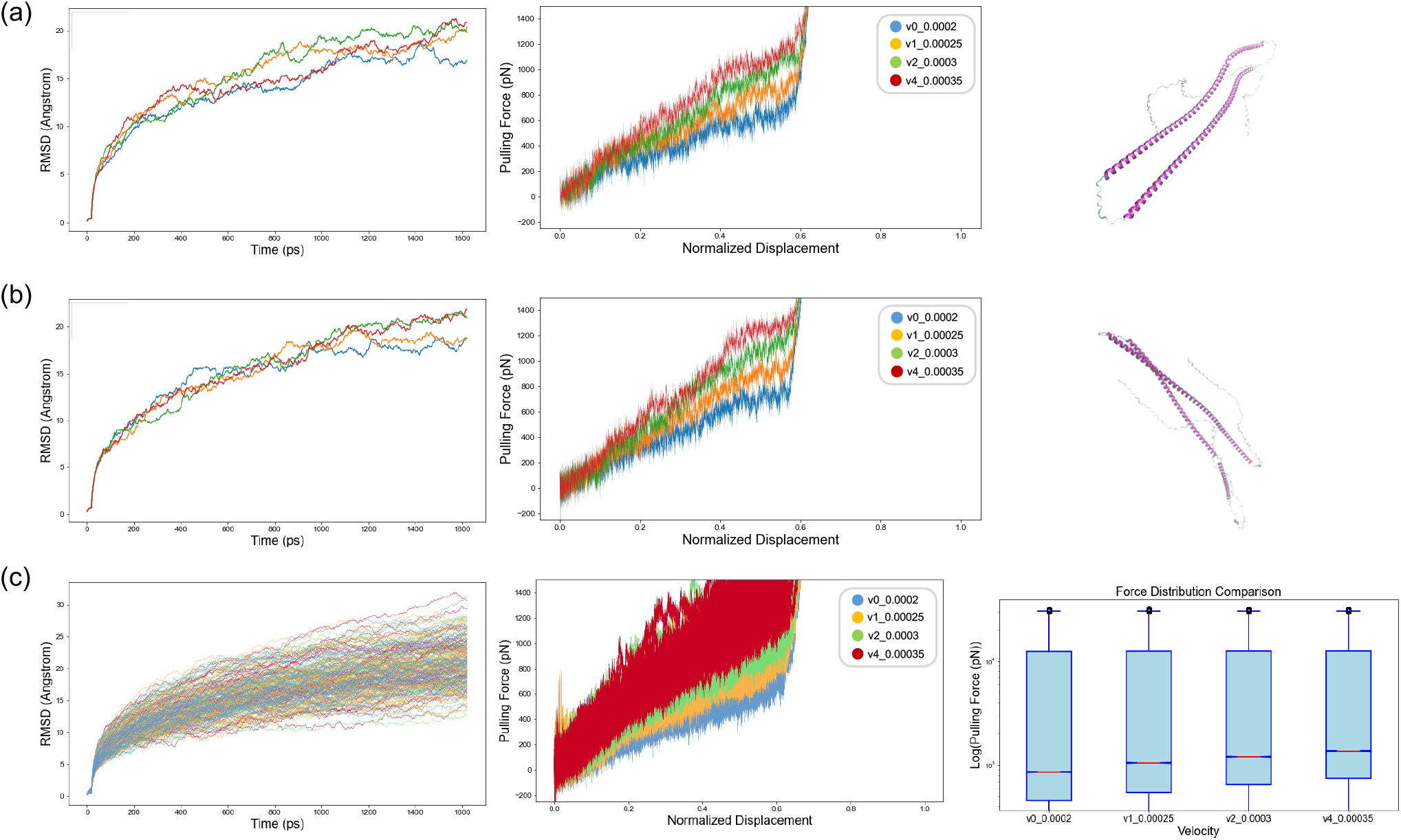
Molecular dynamics simulation performance under controlled SMD pulling velocities. The root mean square deviation (RMSD) and force-displacement plots from molecular dynamics (MD) simulations are presented for (a) KRT35, (b) KRT85, and (c) all 51 keratin proteins collectively. In panels (a) and (b), simulations were performed on Type I (KRT35) and Type II (KRT85) keratin proteins under four different pulling rates (the velocity labels denote pulling increments in Å/timestep, corresponding to 0.1, 0.125, 0.15, and 0.175 Å/ps with a 2 fs timestep). The RMSD plots indicate that both proteins reached stable states during equilibration, as the RMSD values converge across different velocities. The force-displacement plots reveal an almost linear relationship between force and displacement, with higher pulling velocities requiring larger forces to stretch the protein to the same length. Both proteins exhibit helix-turn-helix structures with random coils at the terminal ends. In panel (c), the results from 204 equilibration simulations of all 51 keratin proteins are shown. The RMSD plots confirm stable states across all proteins, while the force-displacement plots demonstrate that force increases with displacement. Pulling-velocity-dependent mechanical behavior is observed, as higher velocities result in larger forces and steeper slopes, indicating increased stiffness. The final plot in panel (c) shows the force distribution using a logarithmic scale for easier comparison. The results demonstrate consistent mechanical performance across all keratin proteins, with smaller force variations observed at higher velocities. Pulling-velocity-dependent behavior is further confirmed, highlighting the increased stiffness and forces at higher pulling velocities.

The unfolding behavior of the keratin proteins follows a well-defined sequence: (1) hydrogen bond rupture, (2) secondary structure uncoiling (e.g., alpha-helices and beta-sheets), and (3) backbone stretching as the protein approaches its contour length [47]. The force-displacement curves in Figure 4(c) show clear pulling-velocity-dependent trends under the accelerated SMD conditions used here. At higher pulling velocities, proteins require larger forces to unfold, reflected in the steeper force-displacement responses. This trend is consistent with reduced molecular relaxation during faster molecular pulling, where unfolding becomes more force-driven and less able to sample alternative relaxation pathways. Thus, the observed stiffening should be interpreted as a relative molecular-scale response under accelerated SMD conditions, rather than as a direct prediction of experimentally accessible hair-fiber strain-rate behavior. Additionally, the force distribution box plot in Figure 4(c) indicates changes in force distributions across pulling velocities, with increasing median force values at higher velocities. These results suggest that accelerated pulling can be used to probe relative unfolding resistance and energy dissipation across keratin proteins.

The simulation outputs highlight how accelerated pulling can be used to probe relative unfolding behavior and nanomechanical differences among keratin proteins. These trends provide insight into molecular mechanisms of unfolding resistance and energy dissipation, while remaining distinct from direct experimental strain-rate comparisons. Sample videos of the SMD simulations for KRT35 and KRT85 are included in the Supplementary Material to further illustrate the unfolding dynamics. As noted in Section 4.2, these rate-dependent trends should be interpreted in the context of accelerated SMD pulling conditions, which emphasize relative unfolding behavior and underlying mechanisms rather than direct reproduction of macroscopic hair deformation rates.

Additionally, secondary structure changes were observed during protein unfolding simulations under different SMD pulling velocities. The secondary structure transition profiles of sample keratin proteins KRT35 and KRT85 (Type I & II) under the same base pulling velocity (v0) are visualized in Figure 5(a). Overall, both protein types exhibit similar unfolding trends, with the primary transition being the conversion of alpha-helices (blue) into random coils (gray). Notably, beta-sheet content remains minimal, which is consistent with the predominantly alpha-helical coiled-coil architecture of native keratin proteins. Furthermore, strain rate dependence was observed in secondary structure changes during unfolding simulations. Although both proteins exhibit similar helix-to-coil transitions under different SMD pulling velocities, as shown in Figure 5(b) (top: KRT35, bottom: KRT85), higher pulling velocities lead to an increased conversion rate from helices to random coils. This suggests that higher unfolding rates reduce structural stability, accelerating the loss of alpha-helical integrity in keratin proteins.

**Figure 5.**
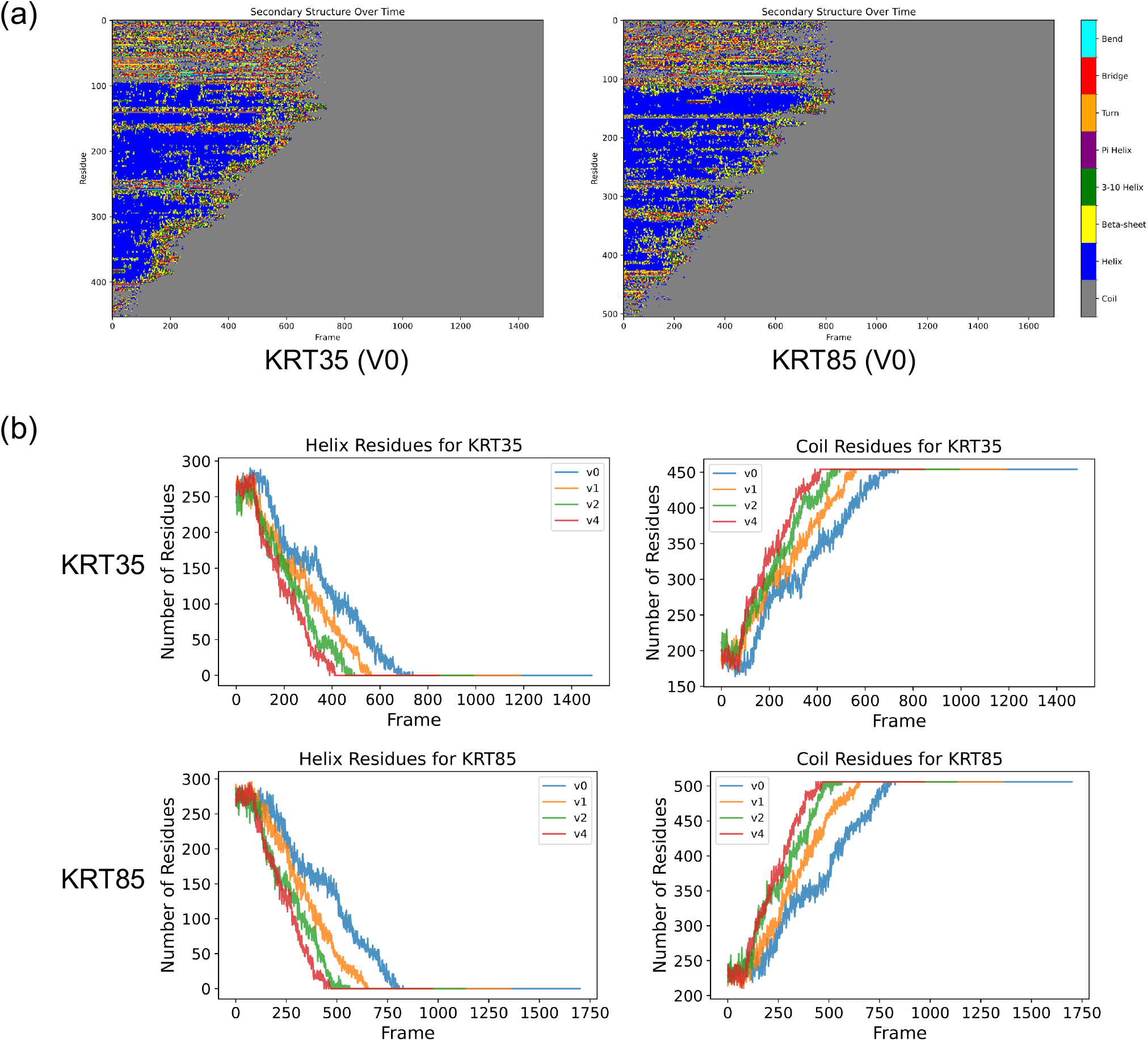
Secondary structure (SS) evolution during protein unfolding. This figure compares the secondary structure transitions of KRT35 (Type I) and KRT85 (Type II) keratin proteins under four controlled SMD pulling velocities. Panel (a): The secondary structure transitions during unfolding are shown for KRT35 and KRT85, as analyzed from the steered molecular dynamics (SMD) simulations. The changes in SS are derived from the trajectory files (.dcd) and analyzed using MDAnalysis and DSSP in Python. The SS transitions are represented by color changes, highlighting the conversion of helices (blue) to random coils (gray). Both proteins show similar trends, with most transitions occurring within the first quarter of the protein unfolding. Panel (b): The specific changes in the primary SS components, helices (left column) and coils (right column), are shown for KRT35 (top row) and KRT85 (bottom row). Both proteins exhibit similar pulling-velocity-dependent behavior, and minimal beta-sheet involvement is observed, which is consistent with the predominantly alpha-helical coiled-coil architecture of keratin proteins. As the pulling velocity increases, the conversion from helices to coils occurs more rapidly, indicating reduced structural stability at higher pulling velocities.

### 2.3 Nanomechanical property characterization and analysis

This section analyzes the nanomechanical properties of keratin proteins, specifically strength and toughness, as characterized from MD simulations at four controlled SMD pulling velocities. The detailed characterization methodology is outlined in Section 4.3.2, and the results are visualized in Figure 6(a). The violin plots in Figure 6(a) reveal bell-shaped distributions, highlighting variability in nanomechanical responses across pulling conditions. Under accelerated SMD pulling, both strength and toughness increase with pulling velocity, consistent with reduced molecular relaxation time and higher resistance to unfolding during faster molecular extension. These trends should be interpreted as relative molecular-scale responses under accelerated SMD conditions, rather than as direct predictions of experimentally accessible hair-fiber strain-rate behavior.

**Figure 6.**
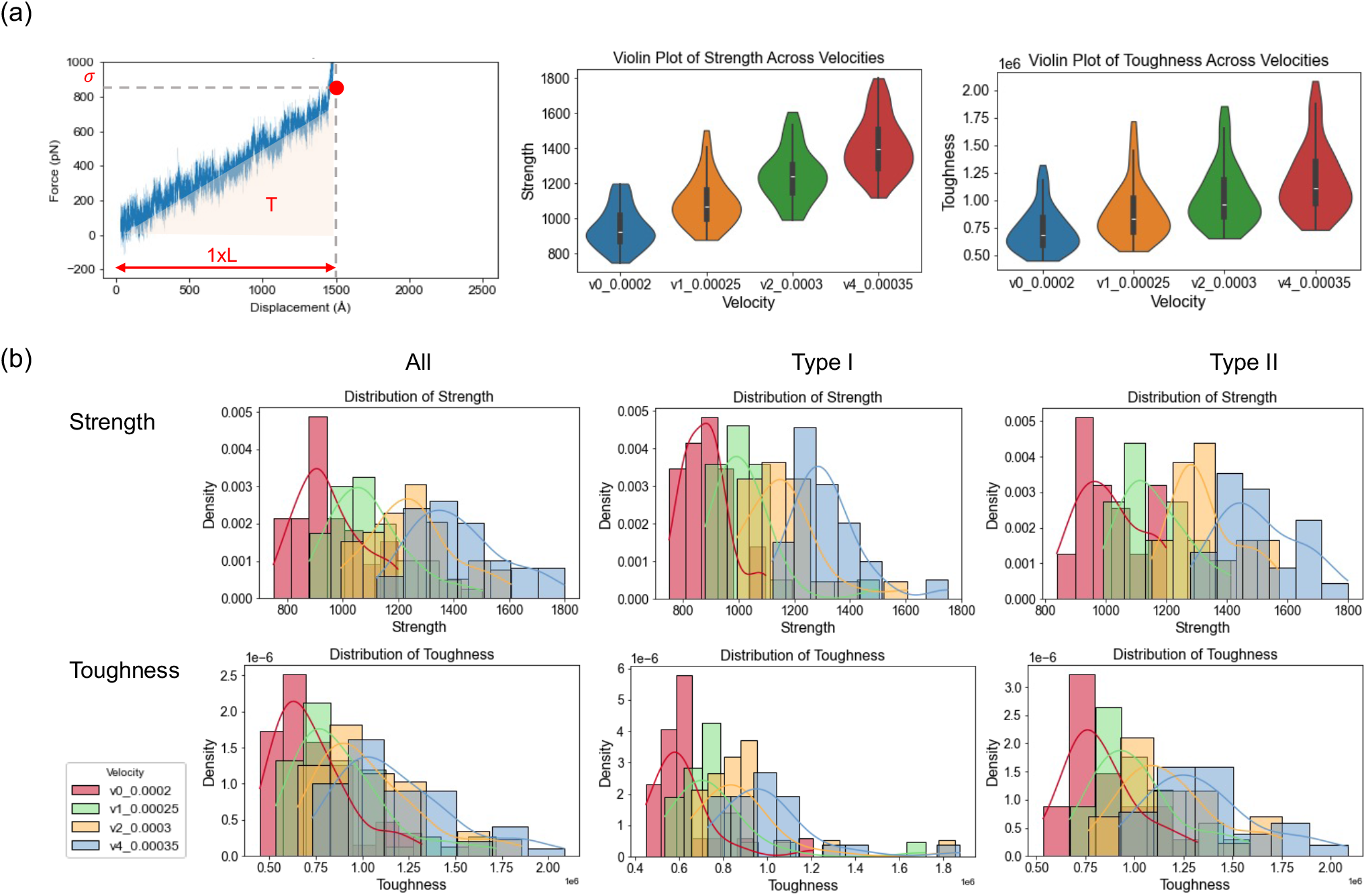
Nanomechanical properties of keratin proteins under controlled SMD pulling velocities. (a) Schematic illustration of nanomechanical property characterization from the force-displacement curve. Strength is defined as the maximum unfolding force (pN), and toughness is defined as the total area under the force-displacement curve (pN *·* Å). The violin plots show that both strength and toughness increase with pulling velocity, indicating pulling-velocity-dependent stiffening and greater energy absorption under faster deformation. (b) Distributions of strength (top row) and toughness (bottom row) across all keratins and stratified by Type I and Type II classes. In all groups, higher pulling velocities shift the distributions toward larger values. Type II keratins generally exhibit higher measured strength and toughness. The higher toughness should be interpreted with the longer average sequence length of Type II in mind, because toughness accumulates over the full extension length.

The distributions of strength and toughness across all proteins and different keratin categories are further illustrated in Figure 6(b). Under the controlled SMD pulling conditions, higher pulling velocities shift the distributions of both strength and toughness toward larger values. This stiffening trend reflects greater resistance to molecular extension and faster energy accumulation within the protein backbone. While the distributions appear approximately normal across the pulling conditions, variations in medians and spreads are evident.

Type II keratin proteins exhibit higher measured strength and toughness values than Type I keratins, as indicated by the shift toward higher values. However, this difference should be interpreted with the effect of molecular size. In this dataset, Type II keratins have a higher average sequence length than Type I keratins, with averages of 544 and 464 residues, respectively. Because toughness is calculated as the area under the force-displacement curve over the full extension length, the higher measured toughness observed for Type II keratins is expected to be strongly influenced by their longer sequence length and larger molecular weight. In contrast, strength is less directly length-dependent. Therefore, the higher strength values may reflect combined effects of molecular size, sequence composition, secondary structure, and unfolding pathways rather than an intrinsically more mechanically robust Type II structure. This interpretation is consistent with the positive correlation between toughness and sequence length shown in Figure 7(b), although this correlation partly reflects the length-dependent definition of toughness. In addition, Type II keratins exhibit a narrower spread while retaining similarly strong strength-toughness coupling, suggesting more uniform unfolding pathways. Type I keratins exhibit greater variability in toughness, indicating a broader range of energy dissipation mechanisms, which may stem from differences in secondary structure flexibility and hydrogen-bonding patterns.

**Figure 7.**
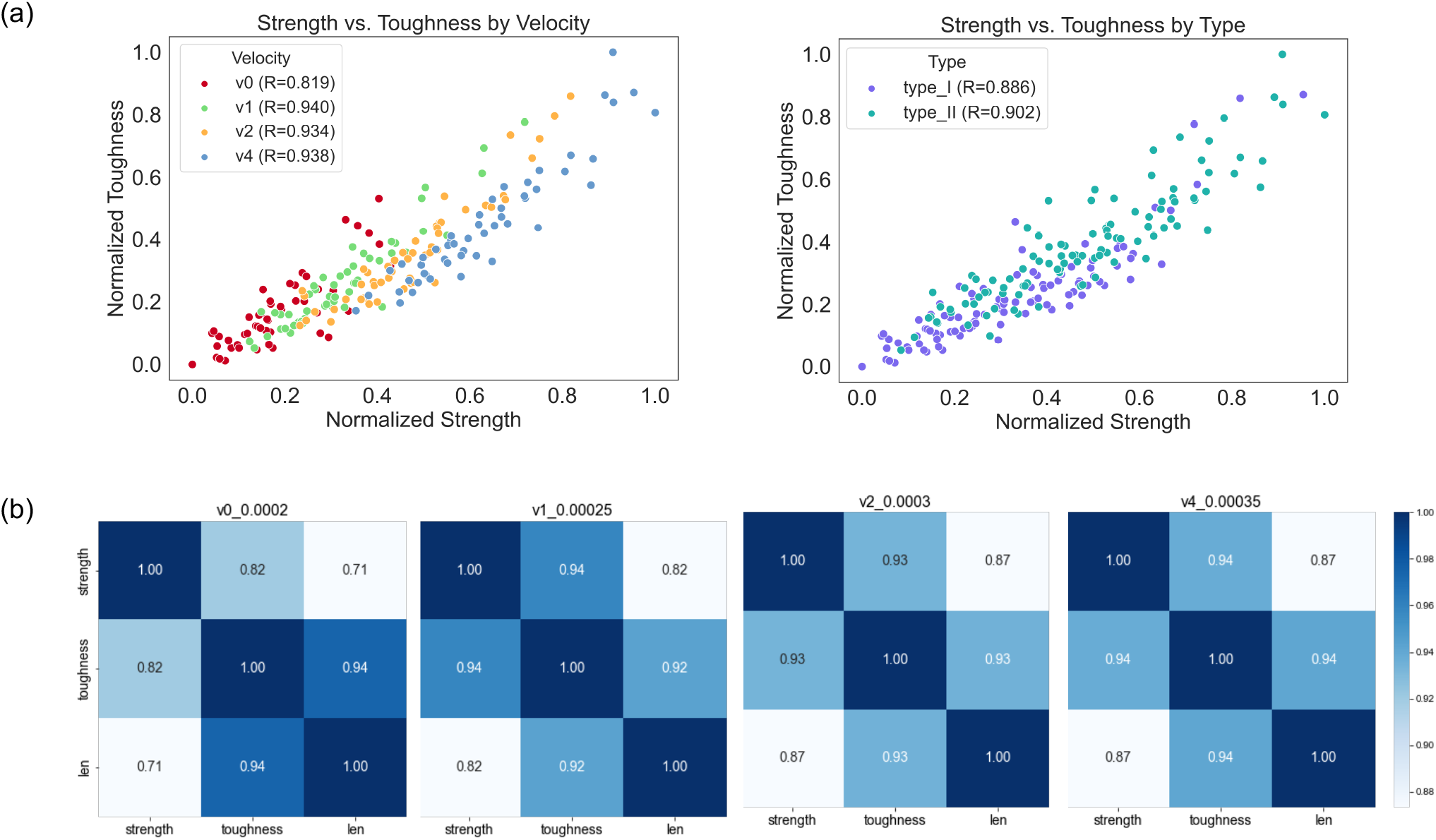
Correlation analysis of nanomechanical properties. (a) Strength-toughness correlations shown using normalized values. The left panel colors the data by pulling velocity and shows that the correlation strengthens at higher pulling velocities (v0: R=0.819, v1: R=0.940, v2: R=0.934, v4: R=0.938). The right panel colors the data by keratin type and shows similarly strong correlations for Type I (R=0.886) and Type II (R=0.902), indicating that the coupling between force resistance and energy absorption is preserved across both keratin classes. (b) Heatmaps of pairwise correlations among strength, toughness, and sequence length at each pulling velocity. These results further show that the strength–toughness relationship remains strong across all velocities, while the correlation between toughness and length becomes more pronounced at higher pulling velocities, supporting the increasing consistency of structure–property coupling under faster deformation.

The updated correlation analysis in Figure 7(a) further refines this interpretation. When grouped by pulling velocity, strength and toughness exhibit strong positive correlations across all velocity groups, with correlation coefficients increasing from *R* = 0.819 at *v*_0_ to approximately *R* ≈ 0.94 at higher pulling velocities. This indicates that, under faster molecular pulling, keratin proteins exhibit more consistent coupling between peak unfolding resistance and total energy dissipation. When stratified by keratin type, both Type I and Type II proteins retain similarly strong correlations (*R* = 0.886 and *R* = 0.902, respectively), indicating that the strength-toughness scaling relationship is largely preserved across both classes. This interpretation is further supported by the per-velocity correlation heatmaps in Figure 7(b), where strength and toughness remain strongly correlated across all pulling velocities, and the correlation between toughness and sequence length becomes more pronounced at higher velocities. This suggests that energy dissipation becomes more systematically linked to molecular size under accelerated pulling, consistent with more uniform unfolding pathways.

Overall, keratin proteins exhibit pulling-velocity-sensitive stiffening under accelerated SMD conditions, where strength and toughness increase at higher pulling velocities. The consistently strong correlation between strength and toughness confirms that mechanically stronger keratins also absorb more energy before unfolding, reinforcing their energy-dissipative role. Additionally, protein length shows a stronger and more consistent positive association with toughness than with strength, which is expected because toughness accumulates force over the full extension length. Thus, molecular size should be considered when interpreting toughness differences across keratin classes. Further analyses of additional mechanical and structural properties, together with regression statistics for the fitted structure-property relationships, are discussed in the following sections and summarized in Table 3. This intrinsic coupling between strength and toughness is further examined through pooled and velocity-specific analysis in Figure 8(b).

**Figure 8.**
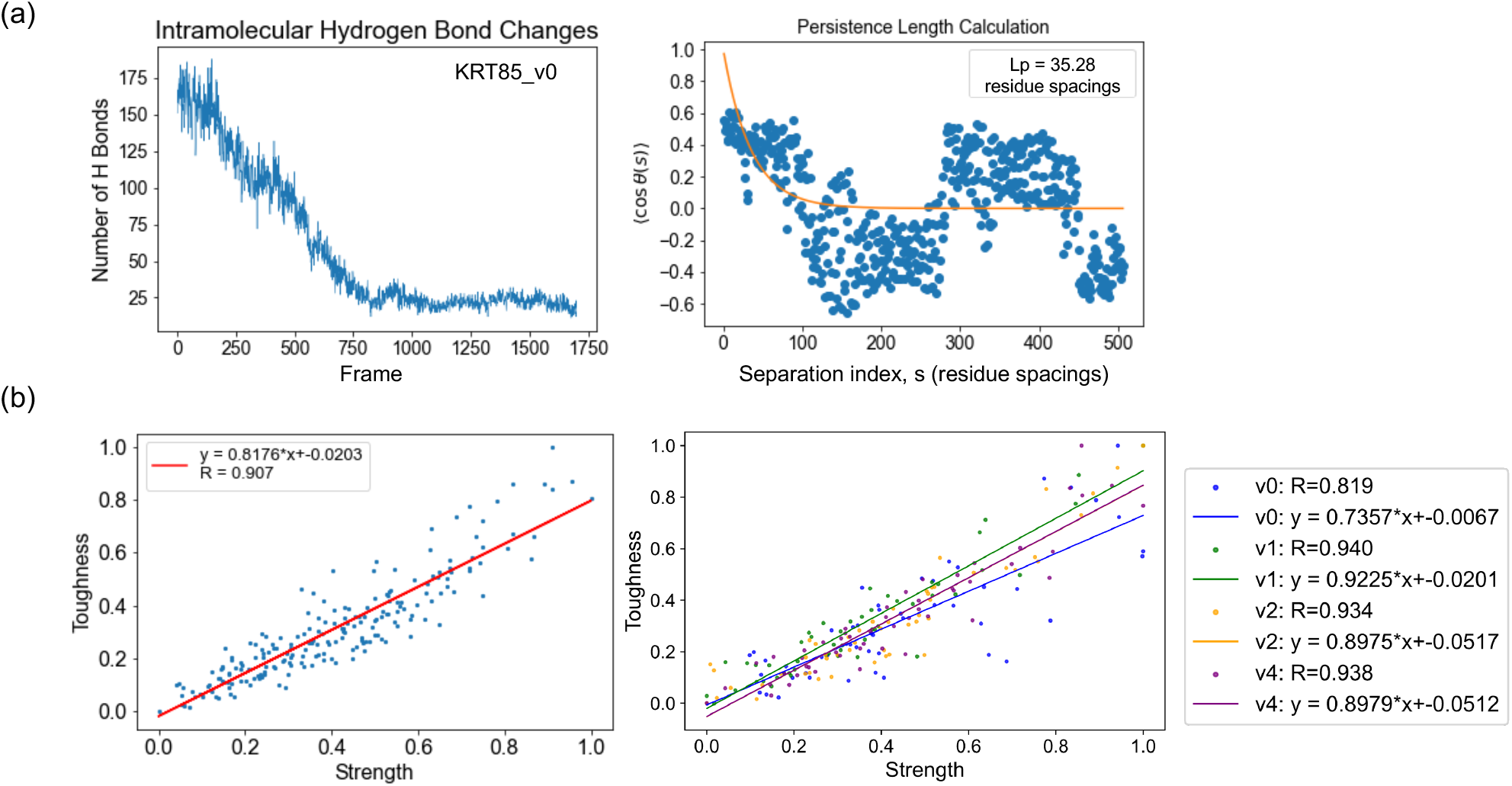
Mechanistic analysis of keratin protein unfolding and strength-toughness coupling. (a) Unfolding-related structural descriptors for representative keratin protein KRT85 at pulling velocity v0. Left: evolution of intramolecular hydrogen bonds during unfolding, showing a progressive decrease in H-bond number as deformation proceeds, consistent with loss of stabilizing interactions and structural destabilization. Right: persistence length (*L*_*p*_) estimation from tangent-tangent correlation decay, obtained by fitting the exponential decay of ⟨cos *θ*(*s*) ⟩ as a function of backbone separation index *s* [58], where *s* is measured in C*α* residue-spacing units. For the KRT85 example, the fitted apparent persistence length is *L*_*p*_ = 35.28 residue spacings, corresponding to approximately 127 Å (12.7 nm) using the 3.6 Å per-residue contour-length conversion. This value should be interpreted as an effective persistence length of the full keratin monomer, including flexible terminal and coil regions, rather than that of an isolated ideal alpha helix. (b) Correlation between normalized strength and normalized toughness. Left: pooled dataset across all pulling velocities, showing a strong positive linear relationship. Right: separate fits for individual velocity-specific groups, showing that strength and toughness remain strongly coupled within each velocity subset. The pooled fit captures the overall global scaling trend across all 204 samples, while the velocity-specific fits reveal rate-dependent differences in the strength-toughness relationship. All fitted linear relationships shown in panel (b) were statistically significant (*p <* 0.001).

### 2.4 Property correlation and scaling

Following the characterization of nanomechanical properties (strength and toughness) from protein simulations, the correlations between mechanical and molecular properties were examined to further investigate the mechanical behavior of keratin proteins under different SMD pulling velocities. The calculation of other molecular properties, including protein sequence properties, secondary structure-related properties, and simulation trajectory properties, is detailed in Sec-tion 4.3.3. In this study, linear property correlations were analyzed to identify statistical relationships between structural features and unfolding behavior, which can provide insights into underlying mechanisms. These correlations were evaluated using a correlation matrix and pairwise correlation plots, with significant relationships summarized in Figure 7, Figure 8(b), and Figure 9. Statistical significance and 95% confidence intervals for the fitted linear relationships in Figure 9 are summarized in Table 3. The key correlations are discussed below, with comparisons across individual pulling-velocity subsets, keratin types, and the combined dataset.

**Figure 9.**
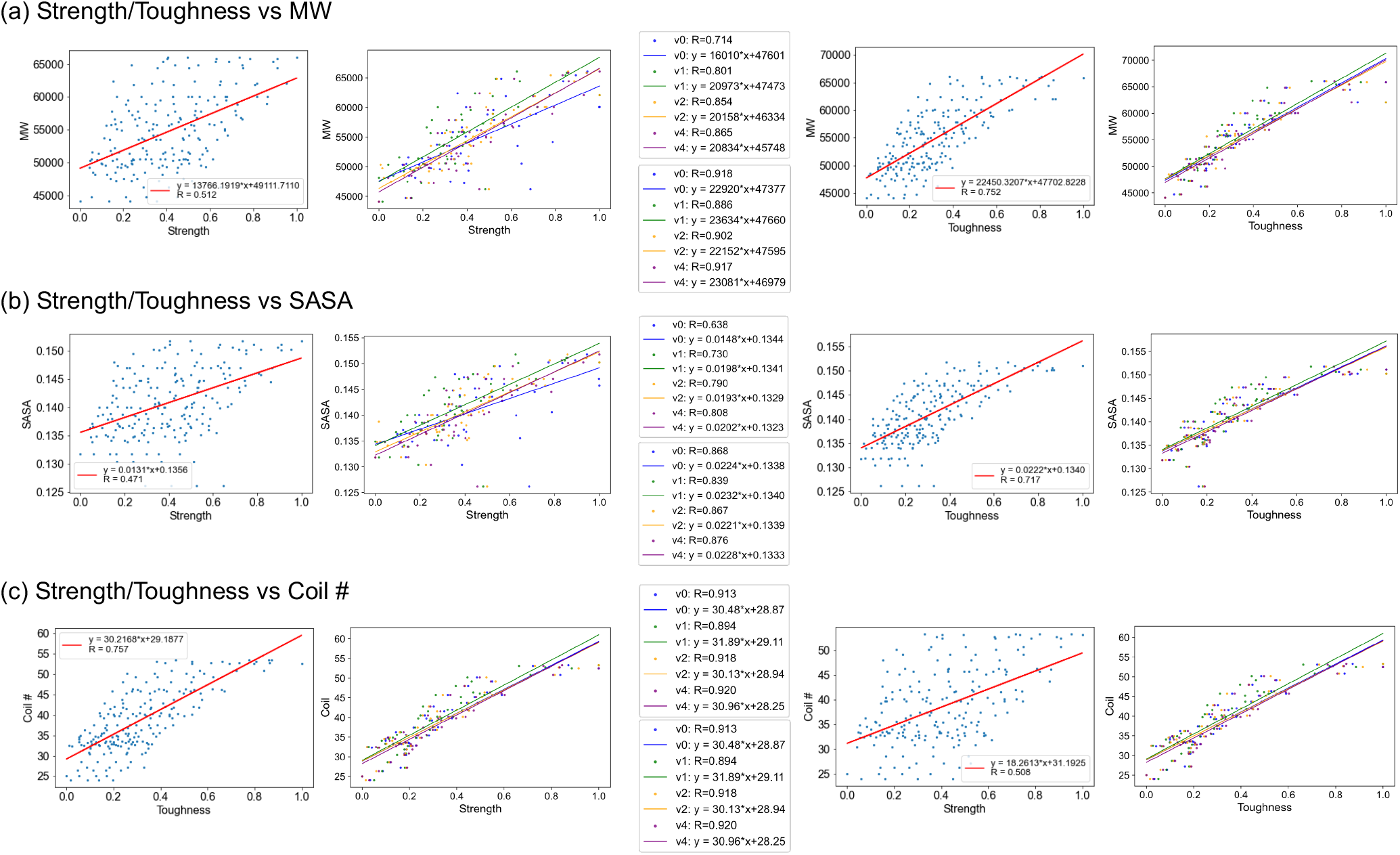
Correlations between nanomechanical properties and molecular descriptors. (a–c) Relationships between strength/toughness and molecular features, including molecular weight (MW), solvent-accessible surface area (SASA), and coil content. Strength and toughness are normalized values; MW is in Da, SASA in nm^2^, and coil content in %. Each row presents correlations for strength (left two panels) and toughness (right two panels), with comparisons between the combined dataset across all velocities and the individual velocity-specific groups. The central annotation boxes summarize the fitted linear relationships, with the upper boxes corresponding to strength and the lower boxes corresponding to toughness, including the regression expressions and correlation coefficients (*R*) for each condition. In general, stronger correlations are observed within individual velocity groups and at higher pulling velocities, indicating more uniform unfolding behavior under faster deformation. All fitted linear relationships shown in this figure were statistically significant (*p <* 0.001). The corresponding regression statistics and 95% confidence intervals of the fitted slopes are summarized in Table 3. While similar trends are observed for both strength and toughness, subsequent mechanistic analysis focuses on toughness, which directly reflects energy dissipation during protein unfolding. Additional multivariable and partial-correlation analyses used to quantify the underlying mechanisms are provided in Table 2, where mechanistically interpretable descriptors are selected for modeling. (a) *Strength/Toughness vs MW:* Higher correlations are observed for individual pulling-velocity subsets than for the combined dataset, likely because distinct velocity-dependent trends are preserved within subsets while variability is mixed in the full dataset. At higher pulling velocities, larger molecular weight is associated with greater mechanical resistance and energy dissipation, consistent with more uniform unfolding and rupture behavior under faster deformation. The toughness-MW relationship should be interpreted partly as a consequence of the length-dependent definition of toughness, because larger proteins have longer contour lengths over which force is integrated. (b) *Strength/Toughness vs SASA:* Similar trends are observed, with stronger correlations for individual pulling-velocity subsets and at higher velocities. More uniform residue exposure at higher pulling velocities strengthens the relationship between SASA and nanomechanical properties, whereas partial or localized unfolding at lower pulling velocities weakens this dependence. (c) *Strength/Toughness vs Coil Number:* Correlations are stronger within individual velocity groups and generally increase with pulling velocity. At lower pulling velocities, local deformation of coil-rich regions can decouple their contribution from the global nanomechanical response, while higher velocities promote more uniform deformation and thus stronger structure–property relationships.

Because the isoelectric-point distribution is bimodal, corresponding to Type I and Type II keratins, stratified analysis was also considered to determine whether the dataset behaves as two mechanically distinct sub-populations. The results indicate that stratification is informative at the level of property distributions (Figure 6(b)), where Type II keratins exhibit higher strength and toughness than Type I keratins. However, the normalized strength-toughness correlations (Figure 7(a)) remain similar across both types, indicating that the dominant scaling relationship between resistance and energy absorption is conserved across keratin classes. This difference in nanomechanical distributions is consistent with the molecular size effect discussed in Section 2.3. In this dataset, Type II keratins have a higher average sequence length than Type I keratins, which enables more stabilizing interactions and greater energy dissipation during unfolding. Thus, while the strength-toughness scaling is preserved across both keratin classes, the upward shift in Type II toughness is strongly influenced by molecular size because toughness scales with the integration length of the force-displacement curve. The strength differences are less directly determined by length and may reflect combined effects of molecular size, sequence composition, secondary structure, and unfolding pathways.

**Table 1.** Summary of the underlying mechanisms associated with linear correlations between nanomechanical properties (strength and toughness) and selected protein descriptors (MW, SASA, and coil content).

| Property | MW | SASA | Coil |
| --- | --- | --- | --- |
| Toughness | Partly reflects length-dependent integration of force over contour length; larger MW also provides longer unfolding pathways for accumulated energy dissipation. | High SASA allows greater flexibility and energy dissipation | Coil regions unfold to absorb energy and delay failure |
| Strength | Less directly length-dependent; may reflect combined effects of molecular size, sequence composition, and stabilizing interactions. | Moderate SASA contributes to external stabilization against forces | Coil content transmits forces to stronger regions; excessive coil content may reduce strength |

**Table 2.** Multivariable regression and partial correlation analysis for toughness prediction across strain rates. For each velocity, a linear regression model was constructed to predict toughness as a function of molecular weight (MW), coil content, hydrogen-bond changes, and secondary-structure transition counts. Toughness is used here as the primary response variable because it reflects the total energy dissipation during protein unfolding and is therefore more directly related to deformation mechanisms than peak strength. SASA was not included in the multivariable model because it is strongly correlated with molecular size and structural descriptors (e.g., coil content) and therefore does not provide an independent mechanistic contribution. The coefficient of determination (*R*^2^) quantifies the fraction of variance in toughness explained by the model. MW coef (p) and Coil coef (p) denote the estimated regression coefficients and their associated p-values in the multivariable model, representing the effect of each descriptor on toughness while controlling for the other included variables. Partial *r* (coil MW) denotes the partial correlation between coil content and toughness after removing the effect of molecular weight, thereby isolating the structural contribution of coil regions beyond protein size. Statistical significance is indica:reted by *(*p <* 0.05), **(*p <* 0.01), and ***(*p <* 0.001).

| Velocity | $R^2$ | MW coef (p) | Coil coef (p) | Partial $r$ (coil \| MW) |
| --- | --- | --- | --- | --- |
| $v_0$ | 0.864 | $2.25 \times 10^{-5}$ (0.008)** | 0.011 (0.088) | 0.344 (0.014)* |
| $v_1$ | 0.817 | $1.50 \times 10^{-5}$ (0.096) | 0.014 (0.042)* | 0.370 (0.008)** |
| $v_2$ | 0.855 | $1.34 \times 10^{-5}$ (0.133) | 0.019 (0.010)** | 0.457 (0.001)*** |
| $v_4$ | 0.869 | $2.11 \times 10^{-5}$ (0.011)* | 0.012 (0.049)* | 0.402 (0.004)** |

**Table 3.** Linear regression statistics for the correlations shown in Figure 9. For the pooled “All” rows, strength and toughness were normalized across all 204 samples together, corresponding to 51 proteins evaluated at four pulling velocities (51 × 4); for velocity-specific rows, normalization was performed within each velocity subset. Slopes are reported with 95% confidence intervals (CI), and the statistic column reports the Pearson correlation coefficient (*R*). All fitted linear relationships listed in this table were statistically significant (*p <* 0.001), where *p* refers to the significance of the fitted linear relationship in the corresponding regression. Strength and toughness were normalized for the regression analysis.

| Velocity | Slope (95% CI)<br>(descriptor unit / normalized property) | Intercept<br>(descriptor unit) | Statistic |
| --- | --- | --- | --- |
| **MW (Da) vs normalized strength** |  |  |  |
| All | 13766 [10563–16970] | 49111.7 | $R = 0.512$ |
| v0 | 16010 [11508–20512] | 47600.7 | $R = 0.714$ |
| v1 | 20973 [16474–25472] | 47472.8 | $R = 0.801$ |
| v2 | 20158 [16633–23682] | 46334.5 | $R = 0.854$ |
| v4 | 20834 [17357–24311] | 45748.5 | $R = 0.864$ |
| **MW (Da) vs normalized toughness** |  |  |  |
| All | 22450 [19723–25177] | 47702.8 | $R = 0.752$ |
| v0 | 22920 [20086–25755] | 47376.7 | $R = 0.919$ |
| v1 | 23634 [20088–27180] | 47660.5 | $R = 0.886$ |
| v2 | 22152 [19114–25190] | 47595.0 | $R = 0.902$ |
| v4 | 23081 [20199–25962] | 46978.6 | $R = 0.917$ |
| **SASA (nm<sup>2</sup>) vs normalized strength** |  |  |  |
| All | 0.013 [0.010–0.017] | 0.136 | $R = 0.471$ |
| v0 | 0.015 [0.010–0.020] | 0.134 | $R = 0.638$ |
| v1 | 0.020 [0.015–0.025] | 0.134 | $R = 0.730$ |
| v2 | 0.019 [0.015–0.024] | 0.133 | $R = 0.790$ |
| v4 | 0.020 [0.016–0.024] | 0.132 | $R = 0.808$ |
| **SASA (nm<sup>2</sup>) vs normalized toughness** |  |  |  |
| All | 0.022 [0.019–0.025] | 0.134 | $R = 0.717$ |
| v0 | 0.022 [0.019–0.026] | 0.134 | $R = 0.868$ |
| v1 | 0.023 [0.019–0.028] | 0.134 | $R = 0.838$ |
| v2 | 0.022 [0.018–0.026] | 0.134 | $R = 0.867$ |
| v4 | 0.023 [0.019–0.027] | 0.133 | $R = 0.876$ |
| **Coil content (%) vs normalized strength** |  |  |  |
| All | 18.3 [14.0–22.6] | 31.19 | $R = 0.508$ |
| v0 | 20.9 [14.7–27.1] | 29.35 | $R = 0.696$ |
| v1 | 28.0 [22.0–34.1] | 28.95 | $R = 0.801$ |
| v2 | 27.0 [22.3–31.7] | 27.41 | $R = 0.854$ |
| v4 | 27.5 [22.7–32.3] | 26.77 | $R = 0.854$ |
| **Coil content (%) vs normalized toughness** |  |  |  |
| All | 30.2 [26.6–33.8] | 29.19 | $R = 0.757$ |
| v0 | 30.5 [26.6–34.4] | 28.87 | $R = 0.913$ |
| v1 | 31.9 [27.3–36.5] | 29.11 | $R = 0.894$ |
| v2 | 30.1 [26.4–33.9] | 28.94 | $R = 0.918$ |
| v4 | 31.0 [27.2–34.8] | 28.25 | $R = 0.920$ |

**Note:** Strength is originally defined in pN and toughness in pN·Å. MW is reported in Da, SASA in nm<sup>2</sup>, and coil content as percentage (%). Slopes are reported in descriptor units per normalized nanomechanical property, and intercepts are reported in the corresponding descriptor units. The velocity labels v0, v1, v2, and v4 correspond to pulling increments of 0.0002, 0.00025, 0.0003, and 0.00035 Å/timestep, respectively, equivalent to 0.1, 0.125, 0.15, and 0.175 Å/ps using a 2 fs timestep.

1. Strength/toughness vs MW: Higher correlations are generally observed for individual pulling-velocity subsets than for the full combined dataset, suggesting that distinct unfolding trends emerge within velocity-specific subsets while pooled analysis introduces additional variability. The toughness-MW correlation should be interpreted cautiously because it partially scales with protein length by construction. In contrast, strength is less directly length-dependent. Therefore, the MW correlations reflect both molecular-size effects and unfolding behavior, rather than purely emergent structure-property relationships.
2. Strength/toughness vs SASA: A similar pattern is observed, with stronger correlations within individual pulling-velocity subsets and, in several cases, at higher pulling velocities. This suggests that under faster loading, residue exposure may be more systematically linked to nanomechanical response, whereas at lower pulling velocities localized unfolding and structural relaxation introduce additional variability.
3. Strength/toughness vs coil content: Correlations are generally stronger within individual pulling-velocity subsets and are often more pronounced at higher pulling velocities. This is consistent with the idea that at lower pulling velocities local deformation of coil-rich regions may occur before full unfolding, whereas at higher velocities coil deformation becomes more uniformly coupled to the global mechanical response.
4. Strength vs toughness: A strong positive correlation is consistently observed across all strain rates and keratin types (Figure 7(a)), indicating a robust coupling between force resistance and energy absorption. This relationship remains strong across both individual pulling-velocity subsets and the combined dataset, reflecting an intrinsic coupling between peak unfolding force and total energy dissipation. While minor variations in correlation strength are observed across velocities, the overall scaling behavior is preserved under all loading conditions.

In summary, for structure-property relationships involving descriptors such as SASA and coil content, local pulling-velocity-specific correlations tend to be stronger than global correlations across the combined dataset, reflecting distinct unfolding behaviors under different pulling conditions. For MW and sequence length, the correlations with toughness should be interpreted with additional caution because toughness is partly length-dependent. Persistence length, however, did not exhibit strong or consistent linear correlations with nanomechanical properties, suggesting that it reflects local chain stiffness rather than serving as a dominant linear predictor of global mechanical response. By comparison, the strength–toughness relationship remains consistently strong across both pooled and velocity-specific analyses, indicating an intrinsic coupling between force resistance and energy dissipation. In addition, the velocity-dependent correlation effects of keratin proteins are observed: at higher velocities, proteins undergo more consistent deformation and unfolding behavior, leading to stronger linear relationships between mechanical and structural properties; whereas at lower pulling velocities, relaxation, plastic deformation, and localized unfolding introduce structural variability, reducing correlation strength. This trend is further supported by the heatmaps in Figure 7(b), where correlations among strength, toughness, and sequence length become more pronounced at higher pulling velocities. The underlying mechanisms governing these relationships are further summarized in Table 1.

Beyond these linear correlations, the increase in toughness with molecular weight should be interpreted as a combination of metric definition and unfolding mechanism. Because toughness is calculated by integrating force over the full extension length, larger proteins with longer contour lengths can accumulate more area under the force-displacement curve even when their average unfolding force is similar. At the same time, longer proteins may provide longer unfolding pathways and more opportunities for energy dissipation through hydrogen-bond rearrangement, secondary-structure uncoiling, and backbone extension. Therefore, the toughness-MW relation-structural properties; whereas at lower pulling velocities, relaxation, plastic deformation, and localized unfolding introduce structural variability, reducing correlation strength. This trend is further supported by the heatmaps in Figure 7(b), where correlations among strength, toughness, and sequence length become more pronounced at higher pulling velocities. The underlying mechanisms governing these relationships are further summarized in Table 1.

Beyond these linear correlations, the increase in toughness with molecular weight should be interpreted as a combination of metric definition and unfolding mechanism. Because toughness is calculated by integrating force over the full extension length, larger proteins with longer contour lengths can accumulate more area under the force-displacement curve even when their average unfolding force is similar. At the same time, longer proteins may provide longer unfolding pathways and more opportunities for energy dissipation through hydrogen-bond rearrangement, secondary-structure uncoiling, and backbone extension. Therefore, the toughness-MW relation-ship reflects both built-in length scaling and molecular deformation mechanisms, rather than a purely emergent structure-property relationship. Coil-rich regions may additionally contribute through entropic elasticity, where configurational freedom enables gradual extension and energy absorption prior to structural failure. In contrast, alpha-helical domains contribute through enthalpic mechanisms, including hydrogen bond rupture and backbone stretching, which require higher forces and contribute to peak resistance. Proteins with larger molecular weight provide longer unfolding pathways, enabling a more sequential progression of these mechanisms and increasing the total accumulated energy dissipation. Therefore, the observed correlations are consistent with contributions from both structural size effects and domain-level unfolding processes. This interpretation is consistent with the unfolding sequence described in Section 2.3, where hydrogen bond rupture, secondary structure uncoiling, and backbone stretching occur progressively during protein extension.

To further quantify the physical mechanisms underlying the observed structure–property relationships, multivariable regression and partial correlation analyses were performed across different SMD pulling velocities (Table 2). While Figure 9 shows similar correlations for both strength and toughness, the following analysis focuses on toughness, as it directly reflects the total energy dissipation during protein unfolding and is therefore more suitable for mechanistic interpretation. For each velocity, a linear regression model was constructed to predict toughness as a function of molecular weight, coil content, hydrogen-bond changes, and secondary-structure transition counts. In this framework, the regression coefficients quantify the independent contribution of each descriptor to toughness while controlling for the others, and the coefficient of determination (*R*^2^) measures the fraction of variance explained by the model. SASA was not included in the multivariable model because it is strongly correlated with molecular size and structural descriptors (e.g., coil content), leading to collinearity that reduces interpretability of regression coefficients, and therefore does not provide an independent mechanistic contribution.

The regression models show consistently high explanatory power (*R*^2^ ≈ 0.82–0.87), indicating that the selected descriptors capture a substantial fraction of the variance in toughness. Among the selected variables, molecular weight and coil content show the most consistent positive associations with toughness within the multivariable framework, although the statistical significance of individual coefficients varies across strain rates. The positive regression coefficients for molecular weight are consistent with both the extensive nature of the toughness metric and longer unfolding pathways that allow greater accumulated energy dissipation. Accordingly, MW should be interpreted partly as a molecular-size factor for toughness rather than only as an independent structural descriptor. Coil content likewise shows a positive contribution, and the partial correlation analysis indicates that coil content remains positively associated with toughness even after controlling for molecular weight, suggesting a structural contribution beyond protein size alone. Because these descriptors are not fully independent, the regression results should be interpreted as statistical associations within the selected feature set rather than strictly independent effects.

These results provide quantitative support for the entropic elasticity mechanism proposed above, in which flexible coil regions enable gradual extension and enhanced energy absorption during deformation. Furthermore, the increasing strength of the coil–toughness relationship at higher pulling velocities suggests that structural features play a more dominant role under conditions of limited molecular relaxation. Hydrogen-bond changes and secondary-structure transition counts were included in the multivariable models, but their effects were weaker and less consistent than those of molecular weight and coil content. This does not imply that hydrogen-bond interactions are unimportant, but rather that their effects are largely coupled with overall structural features and molecular size, and therefore do not emerge as independent predictors of toughness. Overall, these findings are consistent with the trends observed in Figure 9 and provide a quantitative basis for interpreting the underlying deformation mechanisms.

### 2.5 Qualitative comparison with literature

Direct quantitative comparison with experimental tensile data is not possible due to differences in scale, loading conditions, structural hierarchy, and especially the much higher pulling velocities used in SMD [1, 4]. Therefore, the present simulations should not be interpreted as reproducing experimental hair-fiber strain-rate behavior. Nevertheless, the qualitative trend that faster molecular pulling produces higher unfolding resistance is consistent with the broader physical expectation that keratin-based materials exhibit reduced relaxation and increased resistance under faster deformation [1, 57]. In this sense, the simulations provide molecular-scale mechanistic indicators that are compatible with, but not directly equivalent to, experimental observations.

At the molecular level, the dominant helix-to-coil transition observed during unfolding is consistent with the known alpha-helical coiled-coil architecture of keratin proteins. In addition, the strong coupling between strength and toughness and the observed role of molecular length in enhancing energy dissipation are broadly consistent with established structure-property relationships and hierarchical mechanical behavior in keratinous fibrous materials [4, 2]. This is consistent with hierarchical deformation mechanisms in keratin-based fibers, where structural organization across length scales contributes to both force resistance and energy absorption. Although atomistic pulling simulations operate at much higher loading rates and smaller scales than bulk tensile testing, the observed helix-to-coil transitions, strength–toughness coupling, and molecular-size effects suggest that the simulations capture relevant molecular mechanisms for future multiscale models of keratin mechanics [1, 4].

## 3 Conclusions

This study presents a systematic MD framework for comparative characterization of hair keratin unfolding mechanics. To our knowledge, this is among the first dataset-scale molecular simulation studies to compare unfolding behavior, nanomechanical descriptors, and structure–property relationships across a curated set of 51 keratin proteins. A consistent and reproducible workflow was developed, including structure prediction, equilibration, SMD pulling, nanomechanical property characterization, and correlation analysis. The pulling velocities used in this study are accelerated relative to experimentally realizable hair-fiber deformation rates, therefore, the results are interpreted as relative molecular-scale trends and mechanistic indicators rather than direct quantitative predictions of macroscopic strain-rate response. This framework enables the quantification of key nanomechanical properties (strength and toughness) and their correlations with molecular protein features, providing a foundation for future data-driven and machine learning-based predictive models.

Our findings show that, under accelerated SMD pulling conditions, keratin proteins exhibit pulling-velocity-sensitive increases in unfolding force and energy absorption. A strong correlation between strength and toughness confirms that mechanically stronger keratins also absorb more energy during unfolding, highlighting their energy-dissipative role. Additionally, protein length contributes significantly to toughness but has a weaker influence on strength, underscoring the role of molecular size and structural hierarchy in keratin nanomechanics. By examining structure-property correlations, this work shows that local (individual strain-rate) correlations are stronger than global (mixed strain-rate) correlations, indicating that distinct unfolding mechanisms emerge under different deformation rates.

These findings provide a quantitative foundation for comparative keratin unfolding mechanics and support future multiscale modeling of hierarchical hair fibers. By linking molecular structure, controlled pulling response, and nanomechanical descriptors, this work provides mechanistic insights relevant for connecting atomistic simulations to higher-level material behavior. Despite these findings, several limitations remain. The dataset size is limited and may not fully capture the diversity of keratin proteins. The use of implicit solvent modeling neglects explicit solvent effects on unfolding dynamics. In addition, this study does not yet include direct comparison with experimental tensile testing data. Future work will incorporate qualitative and quantitative comparisons with literature-reported keratin and hair fiber mechanics to further validate the simulation results.

Future work will focus on expanding the dataset and exploring non-linear relationships (e.g., Weibull analysis and physics-informed neural networks) to improve predictive capability. For example, while persistence length provides insight into local chain stiffness, its relationship with nanomechanical properties was not found to be strongly linear in the current analysis, suggesting that more complex or non-linear structural descriptors may be required to capture its role in mechanical behavior. A key direction is to bridge sequence-level nanomechanical properties with fiber-level mechanical behavior, enabling multiscale modeling of hierarchical hair fibers and their fracture mechanisms. Furthermore, integrating machine learning approaches for forward prediction and inverse design of keratin proteins offers a promising pathway for optimizing mechanical performance in biomaterials and structural applications.

## 4 Materials and methods

In this section, we detail the materials and methods used in this study. The dataset development process includes keratin protein sequence collection, systematic protein folding, and visualization. The molecular dynamics simulation procedure is described, covering method selection considerations, simulation protocols, and automated pro-cessing for result collection. Finally, we outline various applied analysis procedures, including nanomechanical characterization and other protein property assessments. A detailed analysis and discussion of corresponding results are presented in Section 2.

### 4.1 Dataset development

A hair keratin protein dataset was developed for this study and utilized in molecular dynamics simulations. The dataset consists of 51 folded hair keratin protein structures in PDB format, capturing potential structural uncertainties. All collected data is summarized in a CSV file provided in the supplementary materials, along with the corresponding calculated protein properties. Additionally, the FASTA protein sequence files and the folded PDB protein files are also included in the supplementary materials.

Initially, 51 keratin protein monomer sequences were collected from UniProt [21] in FASTA format. Sequence-based protein properties were calculated using ProtParam as a Python-based tool [22], as shown in Figure 2(a). The calculated properties include sequence length, MW, instability index, and isoelectric point. The instability index, where values below 40 indicate instability, provides insight into the protein’s stability, while the IsoPt represents the pH at which the net charge is zero, distinguishing acidic and basic keratin types.

Using the collected keratin protein sequences, protein structures were predicted and folded using AlphaFold2 [50] and stored in PDB format, as illustrated in Figure 2(b). The molecular structures were subsequently visualized using VMD [52]. Seven representative hair keratin protein structures were selected for further folding performance evaluation, including four Type I proteins (KRT31, KRT32, KRT35, KRT37) and three Type II proteins (KRT81, KRT85, KRT86), as shown in Figure 3. These keratin structures exhibit a consistent architecture, primarily composed of alpha-helices, with end coils and connecting turns. The folding prediction performance of AlphaFold2 was assessed using pLDDT scores, which evaluate the confidence level of structural prediction for each protein data. As shown in the plots of pLDDT per amino acid, the predictions are highly reliable for secondary structure regions (alpha-helices), whereas lower confidence scores are observed in random coil regions at the termini and connecting turns. An average pLDDT value of 73.74 was calculated for the 51 selected folded keratin protein structures, indicating good folding prediction quality suitable for further molecular simulations [53, 54].

### 4.2 Molecular Dynamics (MD) Simulation

Implicit atomistic MD simulations, including equilibration and SMD, were conducted for 51 hair keratin proteins under four different SMD pulling velocities, resulting in a total of 204 MD simulation runs. Nanoscale Molecular Dynamics (NAMD) [59] was employed with the CHARMM force field to model atomic interactions. Specifically, implicit solvent modeling was implemented using the generalized Born implicit solvent (GBIS) model to reduce computational costs, given the relatively large average protein length of 506 residues and the high-throughput nature of the simulations. The choice of atomistic modeling over coarse-grained approaches ensures sufficient resolution for tracking protein unfolding behavior. For SMD, a base pulling velocity of 0.1 Å/ps was selected, balancing simulation stability and computational efficiency. Three additional velocities (0.125, 0.15, and 0.175 Å/ps) were chosen to investigate strain rate effects under the same criteria. In the figure labels, the corresponding pulling incre-ments are reported in Å per timestep as 0.0002, 0.00025, 0.0003, and 0.00035 Å/timestep. With the 2 fs timestep used in the simulations, these correspond to pulling velocities of 0.1, 0.125, 0.15, and 0.175 Å/ps, respectively. A streamlined workflow was developed to enhance simulation efficiency, incorporating automated job script generation, submission, and result collection, following the methodology in [49]. Detailed simulation protocols for both equilibration and SMD are outlined below.

It is important to note that these pulling velocities are significantly higher than deformation rates encountered in real hair fibers, which differ by several orders of magnitude. Such accelerated pulling is standard in SMD to access unfolding within feasible computational timescales [60, 61]. However, under these high-rate conditions, molecular relaxation is limited, and the resulting mechanical response may be dominated by more uniform, force-driven unfolding pathways. Therefore, the results are interpreted in terms of relative trends and underlying mechanisms rather than direct reproduction of macro-scopic deformation rates. In addition, the use of implicit solvent modeling (GBIS), while computationally efficient, does not explicitly capture water-mediated hydrogen bonding and solvent reorganization dynamics that can influence keratin unfolding behavior. In particular, pulling-velocity-dependent solvent relaxation and hydrogen bond rearrangement may affect the unfolding pathways and force response. Therefore, while the current framework captures key structural and mechanical trends, future work incorporating explicit solvent models will be important for further validation and for resolving solvent-coupled deformation mechanisms.

Each protein was equilibrated before SMD to stabilize its configuration. Equilibration was deemed complete once the root mean square deviation (RMSD) of the protein reached convergence. To ensure stability, all proteins underwent 8×10^5^ equilibration steps at a 2 fs time step under controlled temperature and pressure conditions, following an initial minimization phase of 1×10^4^ steps for temperature initialization and relaxation. Output RMSD plots (Figure 4, first column) demonstrate equilibration convergence for a sample type I protein (KRT35) in panel (1), a type II protein (KRT85) in panel (2), and all proteins in panel (c).

SMD simulations were performed on equilibrated proteins by fixing one terminus while applying tensile force to the other, as illustrated in Figure 1(b2). The four pulling velocities (0.1, 0.125, 0.15, and 0.175 Å/ps) were applied until each protein reached its full contour length, estimated as the product of the number of amino acids and an average residue length of 3.6 Å. Pulling forces were recorded every 0.2 ps. The resulting force-displacement data were collected for further analysis of nanomechanical properties. Force-displacement plots were generated to visualize unfolding behavior and assess nanomechanical characteristics, and to facilitate comparison, the displacements were normalized across all proteins. The second column in Figure 4 presents example results for a type I protein (KRT35), a type II protein (KRT85), and the complete dataset in panels (a), (b), and (c), respectively. The following subsections provide further details on characterization and analysis methods. As a result, the simulations emphasize relative trends and comparative behavior across proteins and strain rates, rather than absolute quantitative agreement with experimental mechanical properties.

### 4.3 Simulation performance analysis

Various computational methods and analysis tools were employed to process simulation outputs and characterize the nanomechanical properties of keratin proteins. The analysis focuses on secondary structure transitions, mechanical responses, and correlations between protein properties and unfolding behavior. Detailed methodologies are outlined in this section.

#### 4.3.1 Secondary structure change analysis

The secondary structure evolution during SMD simulations was quantified using DSSP (Define Secondary Structure of Proteins) [62], which analyzes PDB structures and simulation trajectories to track conformational transitions over time. To streamline this analysis, MDAnalysis [63, 64] and Biopython [65] were employed to extract and process structural data. These tools facilitated the computation of secondary structure profiles and their dynamic changes throughout the simulation. The results, visualized in Figure 5, provide insights into the time-dependent unfolding behavior of keratin proteins under different SMD pulling velocities.

#### 4.3.2 Nanomechanical property characterization

With the simulation output, molecular-level mechanical properties of keratin proteins, such as strength (resistance during unfolding) and toughness (total absorbed energy), were quantified, as illustrated in Figure 6(a). The characterization of these nanomechanical properties is specified and calculated with corresponding equations 1 and 2 as follows:

- Strength (*σ*, pN) is defined as the maximum force observed during unfolding
- Toughness (*T*, pN·Å or 10^*™*^22J) is calculated as the total area under the force-displacement curve

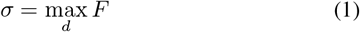

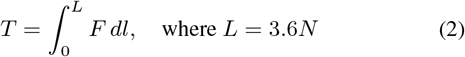

where *d* represents the distance increase during unfolding (Å), *F* is the pulling force (pN), *L* is the contour length (Å), and *N* is the number of amino acids in the targeted protein.

Additionally, to enable quantitative tracking and comparison of force variations, the force vector is stored and resampled to align with the protein sequence length. This is accomplished by averaging force values over segments corresponding to the number of amino acids in each protein. The characterized nanomechanical properties, along with detailed sequence information, are compiled into a CSV file, provided in the supplementary materials. Further analysis is presented in Section 2 and visualized in Figure 7 to 9.

#### 4.3.3 Additional protein property calculation

To analyze the correlation and scaling between protein properties and nanomechanical characteristics, as illustrated in Figure 1(b4), the required protein properties were first computed. In addition to the two nanomechanical properties discussed earlier (strength and toughness), other related properties were categorized into three groups, as summarized below, with their respective calculation methods specified afterward. The correlations between protein sequence, structural, and mechanical properties were then assessed, with notable relationships identified, scaled, and analyzed. The results are summarized and visualized in Figure 7, Figure 8(b) and Figure 9.

- Protein sequence properties: Sequence length, MW, instability index, and isoelectric point.
- Secondary structure properties: Solvent-accessible surface area (SASA), crystallinity, number of coil/helix/sheet structures, and number of transitions between secondary structures along the sequence.
- MD trajectory properties: Persistence length, hydrogen bond (H-bond) changes, and secondary structure transitions.

For protein sequence properties, calculations were performed using the ProtParam tool [22], based on FASTA files containing protein sequences. This systematic computation provided sequence length, MW, II, and IsoPt values, as discussed in Section 4.1.

For secondary structure properties, calculations were conducted using MDTraj[66] and DSSP[65] in Python. Structural features were extracted from PDB data files and simulation trajectories. SASA was computed from equilibration simulations, while the number of helix, sheet, coil structures, and secondary structure transitions were derived from SMD trajectories.

For MD trajectory properties, we quantified H-bond changes and persistence length, two key indicators of protein unfolding stability and mechanical stiffness. H-bond changes, representing stability variations during unfolding, were computed using MDTraj from SMD trajectories. To optimize computational efficiency, the analysis was processed in protein segment chunks. An example of the calculated H-bond change is shown in Figure 8(a). Persistence length, which measures the effective bending stiffness of a polymer chain, was computed using MDAnalysis based on equilibration trajectories. This property was determined by calculating tangent vectors along the protein backbone, computing the cosine of the angle between the initial tangent and subsequent tangent vectors, and fitting an exponential decay function to extract the persistence length [63, 64]. In this calculation, the backbone separation variable was expressed as a C*α* residue-spacing index, when converted to contour distance, one residue spacing was approximated as 3.6 Å, consistent with the contour-length definition used for toughness calculation. An example calculation is illustrated in Figure 8(a). While persistence length provides a measure of effective chain stiffness, subsequent correlation analysis was not found to exhibit strong linear correlations with nanomechanical properties.

Finally, overall property scaling was conducted based on correlation matrix observations and correlation plots linking nanomechanical properties (strength and toughness) to other protein properties. For the pooled correlation analysis across all velocities, strength and toughness were normalized using the full combined dataset of 204 samples, corresponding to 51 proteins evaluated at four strain rates (51 *×* 4). For velocity-specific correlation analyses, normalization was performed separately within each velocity subset before linear fitting. This study primarily focused on linear correlations, while future research may explore nonlinear relationships to further advance the understanding of nanomechanical behaviors.

## Author contributions

M.J.B., W.L., and L’Oreal Team developed the overall concept and the algorithm and oversaw the work. W.L. curated the dataset, conducted MD simulations, analyzed the results, and drafted the paper. W.L. and M.J.B. analyzed the data and edited and wrote the manuscript.

## Data Availability

- Dataset of hair keratin protein in FASTA and PDB format, along with their corresponding protein properties is available as attached as Supplementary Material (‘KRT_property.csv’, ‘PDB_file.zip’and ‘FASTA_file.zip’).
- Dataset of calculated sequence, secondary-structure, MD trajectory, and nanomechanical properties obtained from analysis and simulations under different pulling velocities is available as attached Supplementary Material (‘KRT_mech_seq_SS_MD_properties.csv’).
- Video example of protein simulation, SMD simulation comparison of KRT35 (type I) and KRT85 (type II) under four different SMD pulling velocities.

## Acknowledgments

This work was supported by L’Oreal.

## Codes and datasets

Codes and data are available at: https://github.com/lamm-mit/hair-keratin-mechanics. Additional datasets are available on Hugging Face: lamm-mit/keratin-fasta, lamm-mit/keratin-pdb, and lamm-mit/keratin-mech-seq-ss-md-properties.

